# The Structural Basis for Pacs1-Wdr37 Complex Assembly and Stability

**DOI:** 10.1101/2025.11.05.686565

**Authors:** Le Xiao, Magdalena Grzemska, Xiong Pi, Luming Chen, Michael Turner, Nagesh Peddada, Samantha Calvache, Jessica Jia, Bruce Beutler, Hao Wu, Evan Nair-Gill

## Abstract

Phosphofurin acidic cluster sorting protein 1 (Pacs1) is a multidomain adaptor proposed to bind transmembrane cargo proteins to facilitate their intracellular trafficking. Pacs1 also forms a complex with WD-repeat protein 37 (Wdr37), which is essential for lymphocyte homeostasis. Despite numerous proposed binding partners, a validated structure for Pacs1-containing protein complexes is lacking. Here, we present the cryo-electron microscopy structure of the Pacs1-Wdr37 complex. Pacs1 binds Wdr37 through a conserved interface within its furin-binding region (FBR), the domain previously linked to cargo recognition. This interaction stabilizes Wdr37 and is critical for the expression of both proteins. A gain-of-function mutation in *PACS1* (R203W) causes a highly penetrant neurodevelopmental syndrome. This pathogenic mutation lies on a solvent-exposed surface of the FBR and does not disrupt complex formation. Instead, Pacs1-R203W remains dependent on Wdr37 for stability and its levels can be reduced through targeted Wdr37 proteolytic degradation. Structural homology of the FBR to synaptotagmin C2 domains reveals a previously unrecognized ability of Pacs1 to bind negatively charged phospholipids through a unique positively charged cleft. Together these findings define the structural basis for Pacs1-Wdr37 complex assembly and stability, present potential strategies for Pacs1-mediated neurodevelopmental disease, and suggest novel Pacs1 functions in membrane association.

## INTRODUCTION

Phosphofurin acidic cluster sorting protein 1 (Pacs1) is a multidomain protein previously shown to link phosphorylated acidic clusters in transmembrane cargo proteins to intracellular protein sorting machinery^1^. This model is supported by localization studies and overexpression-based binding assays that indicate Pacs1 controls the distribution of membrane-bound proteins in the secretory and endosomal systems. While the Pacs1 paralog Pacs2 is also predicted to bind phosphorylated acidic clusters, it is thought to operate in distinct trafficking routes with a largely distinct set of client proteins^2^.

Deletion of Pacs1 in mice leads to impaired intracellular calcium (Ca^2+^) release, defective lymphocyte quiescence, and lymphopenia^3^. Pacs2-deficient mice, in contrast, do not display these phenotypes, suggesting that Pacs1 compensates for Pacs2 within the adaptive immune system. Moreover, observed upregulation of Pacs2 within Pacs1-deficient tissues suggests that Pacs2 may be able to compensate for Pacs1 in certain contexts^4^. However, the molecular basis for such compensation is unclear given the anticipated spatial separation of Pacs proteins’ functions within cells.

A recurrent dominant mutation in *PACS1* causes a rare neurodevelopmental syndrome characterized by learning disabilities, craniofacial abnormalities, and seizures^5^. The mutation encodes an R203W substitution within the putative cargo binding domain of Pacs1, commonly referred to as the furin binding region (FBR). A similar syndrome is caused by a recurrent E209K mutation in *PACS2* within its middle region (MR)^6^. Current evidence indicates that these mutations are gain-of-function. Pacs1-R203W is associated with skewed neuronal differentiation *in vitro* and warped neuronal cytoskeletal networks in mouse models^4, 7^. Pacs2-E209K is linked to distorted endoplasmic reticulum-mitochondrial Ca^2+^ signaling^8^. How mutant Pacs proteins mediate these different effects is unclear but is presumed to involve altered interactions with effector proteins.

A limitation in understanding the functions of Pacs proteins, and how disease-associated mutations exert their effects, is the lack of experimentally validated structural information. Our previous genetic studies, together with prior large-scale protein interactome studies, identified WD repeat protein 37 (Wdr37), as a critical Pacs1-binding partner^3, 9^. Wdr37 is a β-propeller-containing adaptor protein with largely unknown functions. Deletion of either Pacs1 or Wdr37 leads to reduced expression of the other^3, 4, 10^. Dominant mutations in *WDR37* also cause a neurodevelopmental syndrome with features overlapping those of *PACS1* and *PACS2* syndromes, suggesting a shared molecular mechanism^11, 12^.

To define the architecture of the Pacs1-Wdr37 complex, we mapped the interaction domains between Pacs1 and Wdr37 and determined the structure of the Pacs1-Wdr37 complex using cryogenic electron microscopy (cryo-EM). The Pacs1 FBR forms a stable interaction with Wdr37, and this interface is fully conserved in Pacs2. The pathogenic Pacs1 R203W mutation does not disrupt complex assembly. Instead, Wdr37 remains essential for mutant Pacs1 expression and can be pharmacologically targeted to reduce pathogenic protein levels. Finally, structural homology with synaptotagmin-like C2 domains revealed that the FBR contains a highly basic cleft capable of binding select acidic membrane phospholipids, thus suggesting a novel function in direct membrane association.

## RESULTS

### The Pacs1 FBR interacts with Wdr37

Pacs1 contains four domains: (1) an N-terminal atrophin-related region (ARR; amino acids [aa] 1-116), named for its sequence similarity to the transcriptional co-activator atrophin; (2) a putative cargo-binding domain (FBR; aa 117-265); (3) a middle region (MR; aa 266-479); and (4) a large C-terminal region (CTR; aa 479-963), which accounts for approximately half of the protein (Fig. 1a). While the FBR is proposed to bind phosphorylated acidic clusters and the MR has been suggested to serve an autoregulatory role, the ARR and CTR lack defined functions^2^.

**Figure 1.**
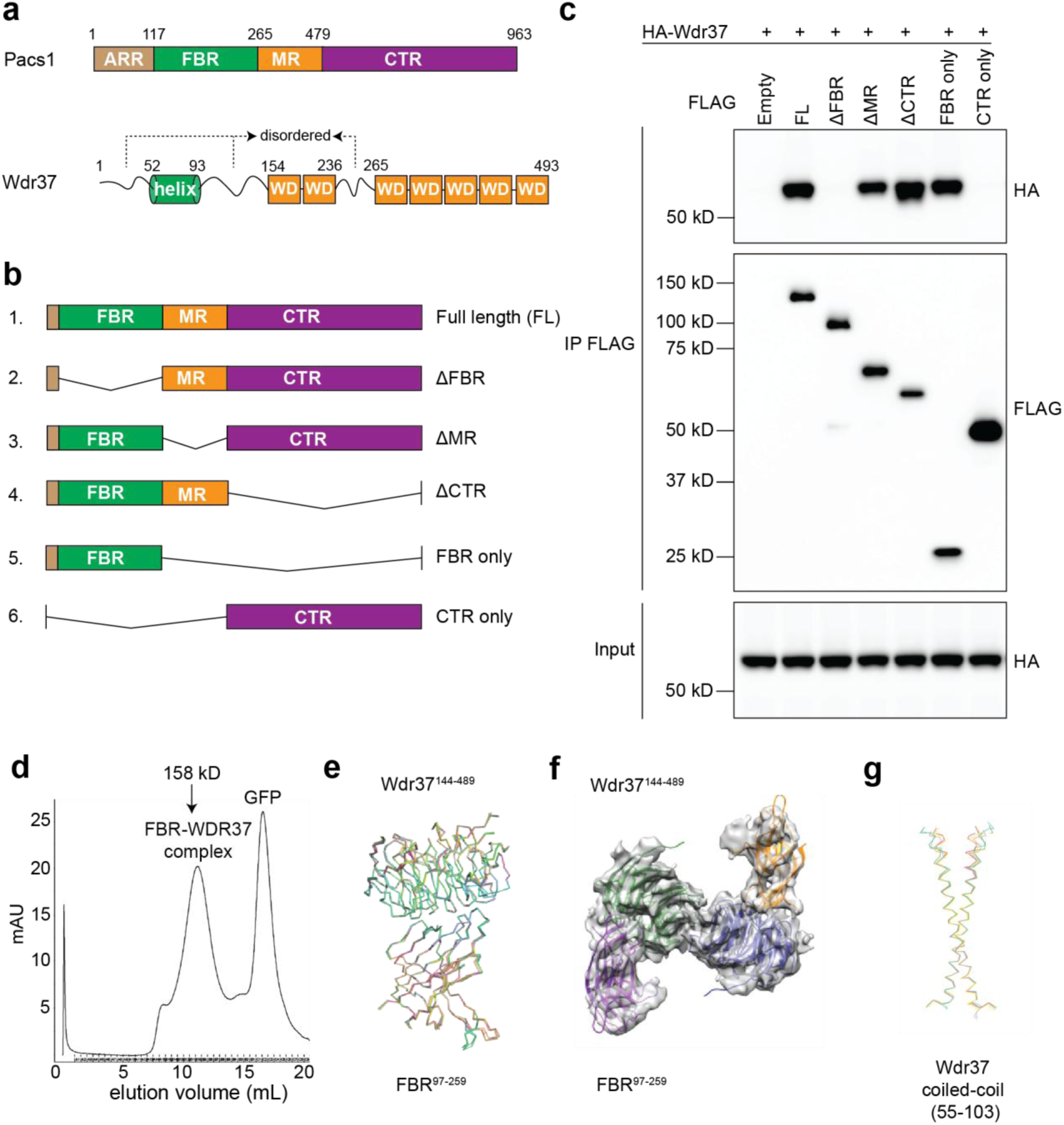
Domain mapping of the Pacs1-Wdr37 interaction. **a** Schematic of Pacs1 and Wdr37 domains. **b** Schematic of Pacs1 constructs used to identify regions required for Wdr37 binding. **c** Co-immunoprecipitation of HA-tagged Wdr37 with FLAG-tagged Pacs1 domain deletion mutants. **d** Size-exclusion chromatography (SEC) profile of purified FBR-Wdr37 complex. SDS-PAGE analysis of SEC fractions corresponding to the FBR-Wdr37 peak. **e** Overlay of five predicted models of FBR-Wdr37 complex from AlphaFold3. **f** The refined FBR-Wdr37 model fit into the cryo-EM density. **g** Overlay of the five predicted models of the Wdr37 coiled-coil domain dimer from AlphaFold3.

Prior circular dichroism analysis predicted that the ARR and MR are intrinsically disordered, whereas the FBR and CTR possess secondary structure^1^. Our initial attempts to purify full-length Pacs1 in complex with Wdr37 from mammalian cells failed to yield suitable particles for cryo-EM analysis, likely due to conformational heterogeneity introduced by the disordered ARR and MR.

To identify the Wdr37-binding region, we performed a domain truncation analysis of Pacs1 (Fig. 1b). Based on our previous findings that the ARR is dispensable for Wdr37 binding, and its absence in Pacs2, we excluded this region from further analysis^3^. We deleted the FBR, MR, and CTR from epitope-tagged Pacs1 and assessed Wdr37 binding by co-immunoprecipitation (co-IP). Deletion of the FBR completely blocked binding to Wdr37 while deletion of either the MR or CTR had no effect. Additionally, co-expression of the FBR alone was sufficient to pull down Wdr37 from cell lysates (Fig. 1c). Thus, the FBR mediates the interaction between Pacs1 and Wdr37. This finding aligns with recent studies in C. elegans showing that the FBR is essential for Pacs1 protein stability^10^.

### Cryo-EM structure determination of the FBR-Wdr37 complex

For single-particle cryo-EM analysis, we co-expressed human FLAG-GFP-FBR (amino acids 91-235) and full-length, strep-tagged human Wdr37 in Expi293F cells followed by lysis under detergent-free conditions. The complex was purified by anti-FLAG immunoprecipitation, followed by cleavage of the FLAG and GFP tags using tobacco etch virus (TEV) protease. The resulting eluate was further purified by size-exclusion chromatography, during which the FBR-Wdr37 complex eluted as a single, symmetric peak (Fig. 1d). SDS-PAGE analysis confirmed that both proteins were present in approximately equimolar amounts, consistent with a 1:1 stoichiometric complex (Fig. 1d).

We imaged purified FBR-Wdr37 complex using cryo-EM but observed severe orientation preference. We thus collected all data at a stage tilt angle of 37 ° to mitigate the problem. Three-dimensional (3D) classification revealed at least two main states (Fig. S1a). 3D refinement resulted in structures of the FBR-Wdr37 complex as a dimer with C2 symmetry and as a monomer, both at 4.5 Å resolution (Fig. S1b-e). We then predicted a 1:1 complex structure with AlphaFold3^13^, which generated five models that were essentially of the same structure (Fig. 1e). The top-ranked model had an interface predicted template modelling (iPTM) score of 0.899 and a predicted PTM score of 0.73, suggesting high confidence. The model fit well with the monomer density, and each half of the dimer density, despite the modest resolutions (Fig. 1f, Table S1).

AlphaFold did not predict the FBR-Wdr37 dimer complex, especially for the Wdr37-Wdr37 interface, which was thus entirely experimentally derived. However, it predicted that a helix spanning aa 55-102 in the Wdr37 N-terminal region could form a coiled coil (CC) dimer (Fig. 1g). This region is not visible in the cryo-EM map, likely due to the existence of a flexible linker (residues 103-144) between the CC and the WD40 domain in Wdr37. The final models contain residues 144-489 of Wdr37 and residues 97-259 of the FBR of Pacs1.

### Overall structure of the FBR-Wdr37 complex

Cryo-EM structure determination revealed a tetrameric complex composed of two Wdr37-FBR heterodimers joined via a Wdr37-Wdr37 interface (Fig. 2a, b). Wdr37 adopts a canonical seven-bladed WD40 β-propeller fold with a central pore. A flexible N-terminus (amino acids 1-154) and a loop between blades 2 and 3 (amino acids 236-265) were not resolved in the structure, likely due to conformational heterogeneity. Within the complex, Wdr37 subunits were arranged such that blades 1-3 (Side A) of each subunit faced one side of the complex, while blades 5-7 (Side B) faced the opposite side (Fig. 2c). The juxtaposed blade 4 regions of each Wdr37 subunit formed a dimer interface and were tilted at a ∼75° angle (Fig. 2b).

**Figure 2.**
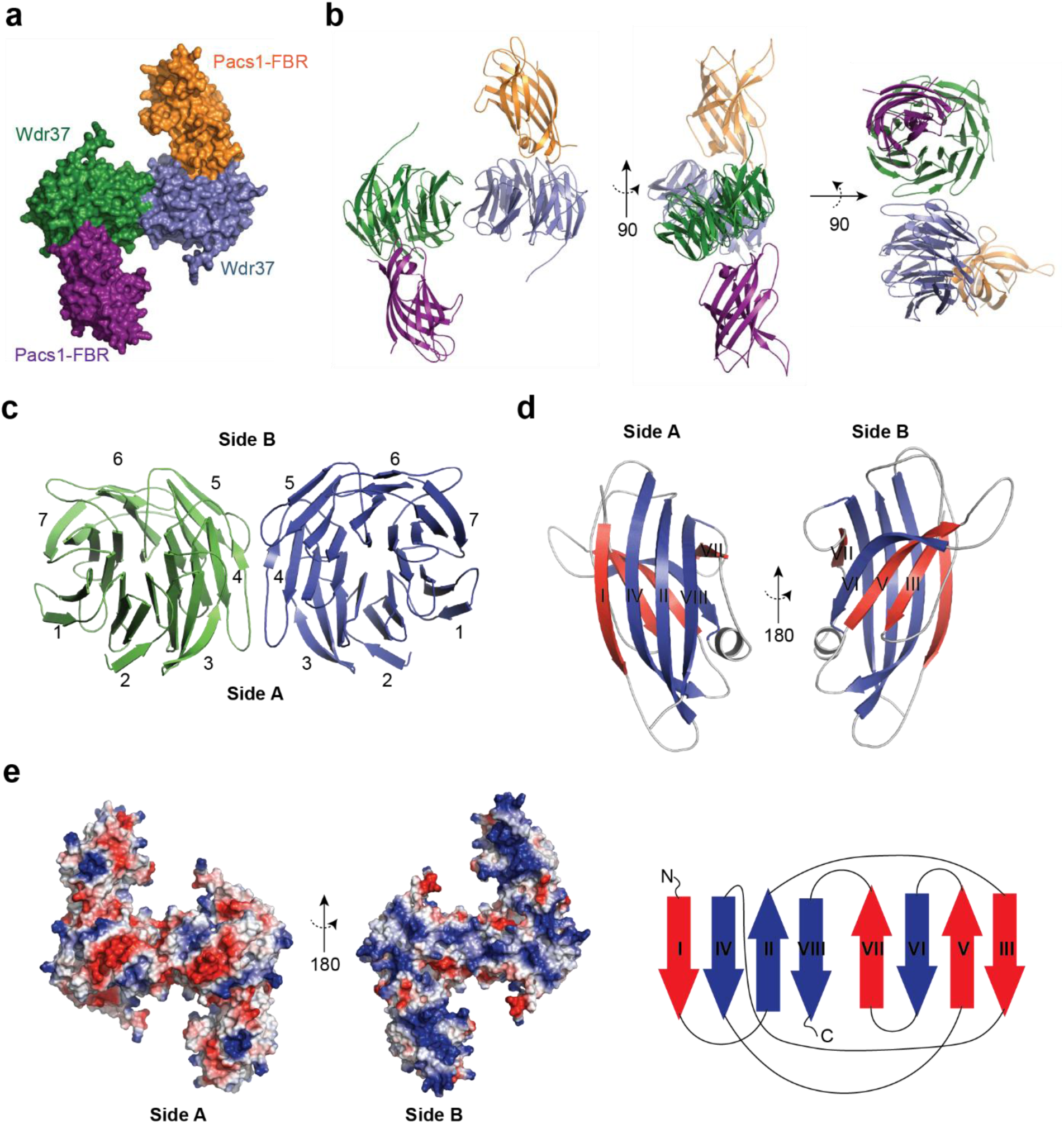
Structural organization of the Pacs1-Wdr37 complex. **a** Cryo-EM density map of the Pacs1-FBR-Wdr37 tetramer. **b** Ribbon models of the complex, shown in orthogonal orientations rotated around the y- and z-axes. **c** Domain schematic of Wdr37 and orientation of β-propeller subunits within the tetramer. **d** β-strand topology of the Pacs1 FBR. **e** Surface electrostatic potential map of Sides A and B of the tetramer.

The FBR adopted an 8-stranded β-sandwich fold. Strands βI, βII, βIV, and βVIII (Side A) formed a convex surface, while strands βIII, βV, βVI, and βVII (Side B) formed a concave surface (Fig. 2d). The FBR exhibited an unusual topology, with βIV inserting between βI and βII in a parallel orientation relative to βI and anti-parallel to βII. The remaining strands were arranged in an anti-parallel fashion. Each FBR subunit was bound to the outer surface of a Wdr37 subunit, distal from the Wdr37-Wdr37 interface, with its concave surface (Side B) facing toward the same side of the tetramer (Fig. 2b).

The overall architecture of the complex displayed twofold rotational symmetry in the x-y plane. However, the surface electrostatic potential was asymmetric when viewed along the z-axis. Side B, which exposed the FBR concave surfaces and Wdr37 blades 5-7, was enriched in basic residues, while Side A showed a predominance of acidic residues (Fig. 2e).

### Mutational Analysis of the Pacs1-Wdr37 Binding Interface

Analysis of the FBR-Wdr37 interface revealed two major contact sites (Fig. 3a). First, the side chain of Pacs1 K184, located in a loop between βIV and βV of the FBR, formed salt bridges with Wdr37 residues D386 and D428, positioned on blades 5 and 6, respectively. Second, the side chain of Pacs1 Q223, located at the end of a short helical segment between βVI and βVII, formed hydrogen bonds with Wdr37 residue N203 on blade 2. Mutating either K184 or Q223 to alanine substantially reduced FBR-Wdr37 binding in co-immunoprecipitation assays (Fig. 3b). Similarly, alanine substitutions at Wdr37 residues D386, D428, or N203 also disrupted complex formation (Fig. 3c, d).

**Figure 3.**
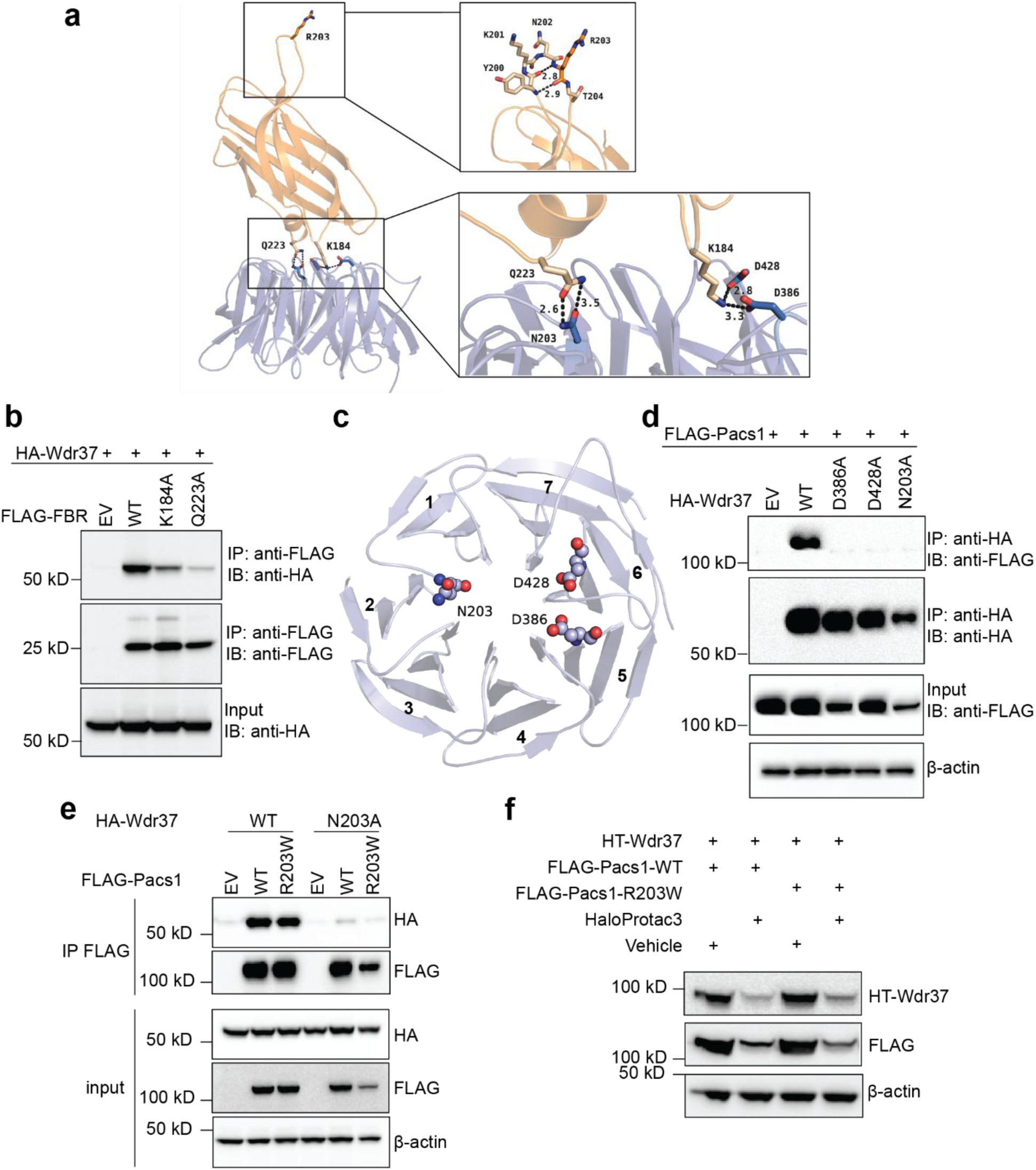
Mutational analysis of the Pacs1-Wdr37 interface. **a** Ribbon representation of the FBR-Wdr37 heterodimer with contact residues highlighted. Structural location of R203 within the FBR-Wdr37 heterodimer. **b** Co-IP of HA-tagged Wdr37 with alanine mutants of Pacs1-FBR at predicted interface residues. **c** Structural location of Wdr37 residues contacting the FBR, mapped onto the β-propeller blades. **d** Co-IP of FLAG-tagged Pacs1 with Wdr37 alanine mutants to disrupt Pacs1 binding. **e** Size-exclusion chromatogram of Wdr37 expressed in the absence of Pacs1-FBR, showing aggregation in the high molecular weight range. **f** Co-IP of Wdr37 with Pacs1-R203W and effect of the Wdr37 N203A mutation. **g** HaloProtac3-mediated degradation of HaloTag-Wdr37 in HEK293T and its effect on Pacs1-WT and Pacs1-R203W protein levels.

Notably, K184 and Q223 are conserved in Pacs2, consistent with the high sequence identity between the FBRs of Pacs1 and Pacs2 and with proteomic evidence that both can bind Wdr37^9^. These contact residues are also conserved in distant orthologs of Pacs1 and Wdr37, supporting a strong functional importance of this interaction (Fig. S2a, b).

When expressed alone, Wdr37 eluted from size-exclusion chromatography in high-molecular weight fractions corresponding to >600 kDa, consistent with the formation of large aggregates (Fig. S3a, b). This behavior is consistent with prior reports that WD40 β-propellers are prone to misfolding and aggregation when not properly stabilized^14, 15^. Expression of Wdr37 mutants that disrupt binding to the Pacs1 FBR also led to reduced Pacs1 levels, mirroring the reciprocal destabilization observed in *Pacs1* and *Wdr37* knockout mouse and nematode models (Fig. 3d)^3, 10^. Together, these results indicate that the Pacs1 FBR plays a key role in stabilizing the nascent Wdr37 β-propeller and support a model in which the Pacs1-Wdr37 complex assembles co-translationally^16^.

### Targeted degradation of Wdr37 reduces mutant Pacs1-R203W

A recurrent pathogenic R203W mutation in *PACS1* causes a highly penetrant neurodevelopmental syndrome^5^. R203 is located within a loop on the distal face of the FBR β-sandwich, opposite the Wdr37-binding interface (Fig. 3a). The peptide backbone of R203 forms hydrogen bonds with neighboring residue Y200, but the R203 side chain is solvent-exposed and does not participate in Wdr37 interaction. Accordingly, the R203W mutation did not disrupt binding to Wdr37 in co-IP assays. Furthermore, mutation of Wdr37 residue N203, which is essential for FBR binding, also blocked interaction with Pacs1-R203W (Fig. 3e). These findings support a model in which R203W preserves core complex assembly but may alter downstream interactions or localization.

Emerging genetic and cellular evidence indicates that R203W acts in a gain-of-function manner^4, 7, 10^. We reasoned that if the mutant protein depends on Wdr37 for stability, then degrading Wdr37 could reduce R203W levels. To test this concept, we employed the HaloTag (HT) system, in which HaloTag-fusion proteins can be targeted for proteasomal degradation upon treatment with the HaloPROTAC3 (HP3) ligand^17^. We co-expressed N-terminal HT-Wdr37 with either FLAG-Pacs1-WT or FLAG-Pacs1-R203W in HEK293 cells. Treatment with HP3 induced rapid degradation of HT-Wdr37 and led to a substantial reduction in both WT and R203W Pacs1 levels (Fig. 3f). These results suggest that targeting Wdr37 for degradation may provide a strategy to reduce levels of pathogenic Pacs1 protein.

### Dimerization and Pacs1 binding are separable features of Wdr37

Cryo-EM analysis revealed that the Pacs1-Wdr37 complex is a tetramer composed of two Wdr37-FBR heterodimers joined through a Wdr37-Wdr37 interface. This interface involves a short β-strand segment within blade 4 (residues H358-Q364) of each Wdr37 subunit, forming a parallel β-sheet at the core of the dimer (Fig. 4a). However, structural data did not resolve the flexible N-terminal region of Wdr37 (residues 1-154), which AlphaFold predicted to form a short coiled-coil (CC) helix between residues 55-103 (Fig. 1g). We hypothesized that this region may also contribute to Wdr37 self-association.

**Figure 4.**
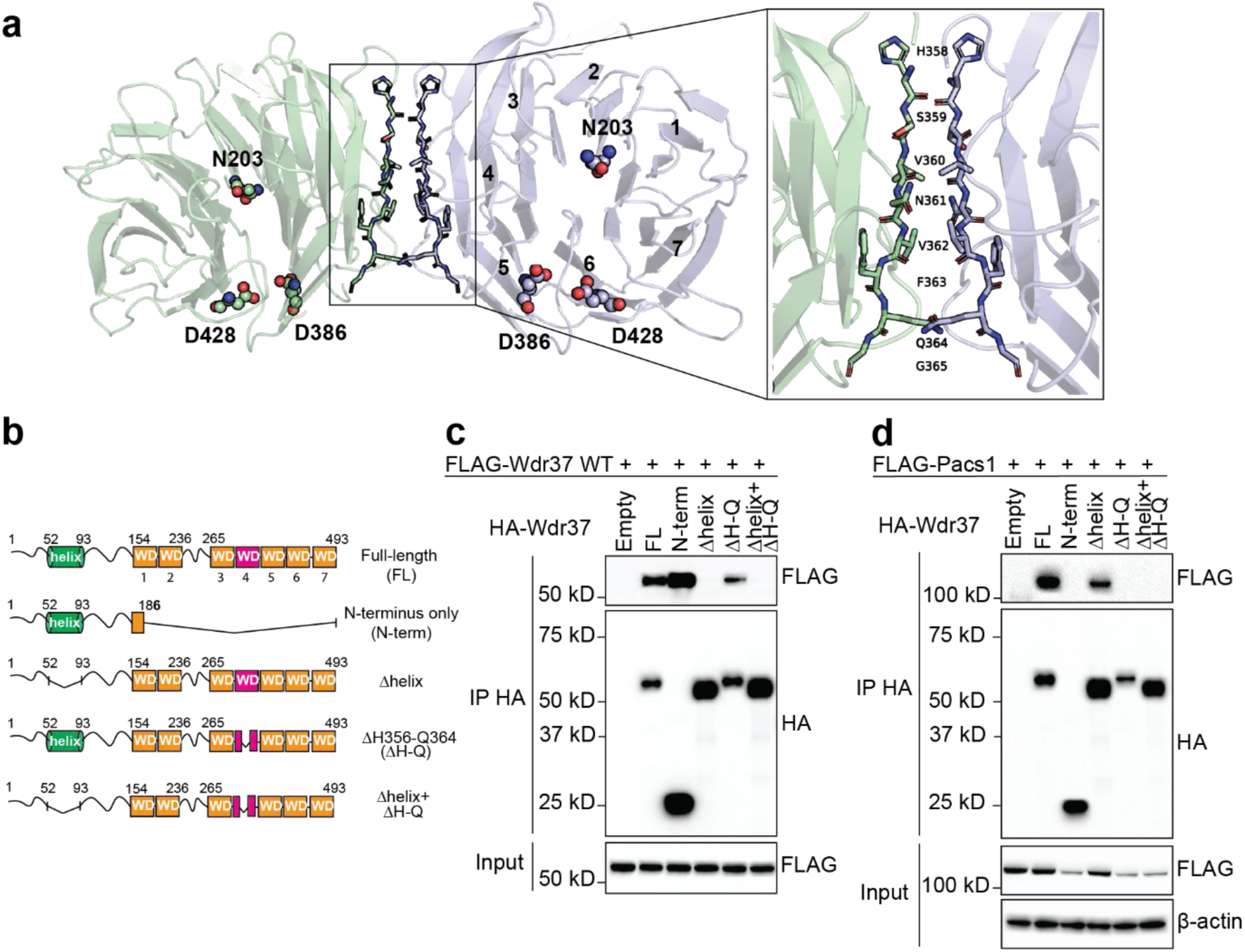
Wdr37 dimerization is mediated by an N-terminal helix. **a** Cryo-EM model of the Wdr37-Wdr37 interface showing β-strand pairing between blade 4 of each subunit. **b** Schematic of Wdr37 domain truncation constructs used to assess dimerization. **c** Co-IP of FLAG-tagged Wdr37 with HA-tagged Wdr37 deletion variants. **d** Co-IP of FLAG-tagged Pacs1 with HA-tagged Wdr37 deletion variants.

To test the contributions of these separate regions to Wdr37 dimer formation, we generated HA-tagged Wdr37 variants with targeted deletions and assessed their ability to co-IP full-length FLAG-tagged Wdr37 (Fig. 4b). A construct containing only the N-terminal region (residues 1-154) was sufficient to pull down full-length Wdr37, while deletion of the predicted CC region (ΔCC) abolished the interaction (Fig. 4c). In contrast, deletion of the structured β-strand region (ΔH-Q; residues 356-364) did not impair dimerization. Therefore, the N-terminal CC is both necessary and sufficient for Wdr37-Wdr37 association.

We next asked whether Wdr37 dimerization is required for Pacs1 binding. Co-immunoprecipitation of Pacs1 with the Wdr37-ΔCC variant revealed that loss of dimerization does not impair Pacs1 binding, indicating that Wdr37 dimerization is not a prerequisite for complex formation (Fig. 4d). However, while the ΔH-Q mutant retained the ability to dimerize, it was unable to bind Pacs1. To dissect this further, we introduced point mutations within the H356-Q364 region and found that substitutions introducing charged residues, particularly V360K and F363E, reduced Pacs1 binding (Fig. S4). These data suggest that this β-strand region, though not strictly required for dimerization, is important for maintaining the structural scaffold of Wdr37 necessary for FBR engagement.

### The Pacs1 FBR shares structural features with phospholipid-binding domains and interacts with negatively charged phospholipids

A Dali structural homology search of the Pacs1 FBR revealed significant similarity to the C2B domain of synaptotagmin (Z-score 11.9; RMSD 2.4 Å, Fig. 5a)^18^. C2 domains are shared across diverse protein families where they mediate binding to membrane phospholipids. Although classical C2 domains form completely anti-parallel β-sandwiches, the Pacs1 FBR adopts an atypical topology with a mix of parallel and anti-parallel strands.

**Figure 5.**
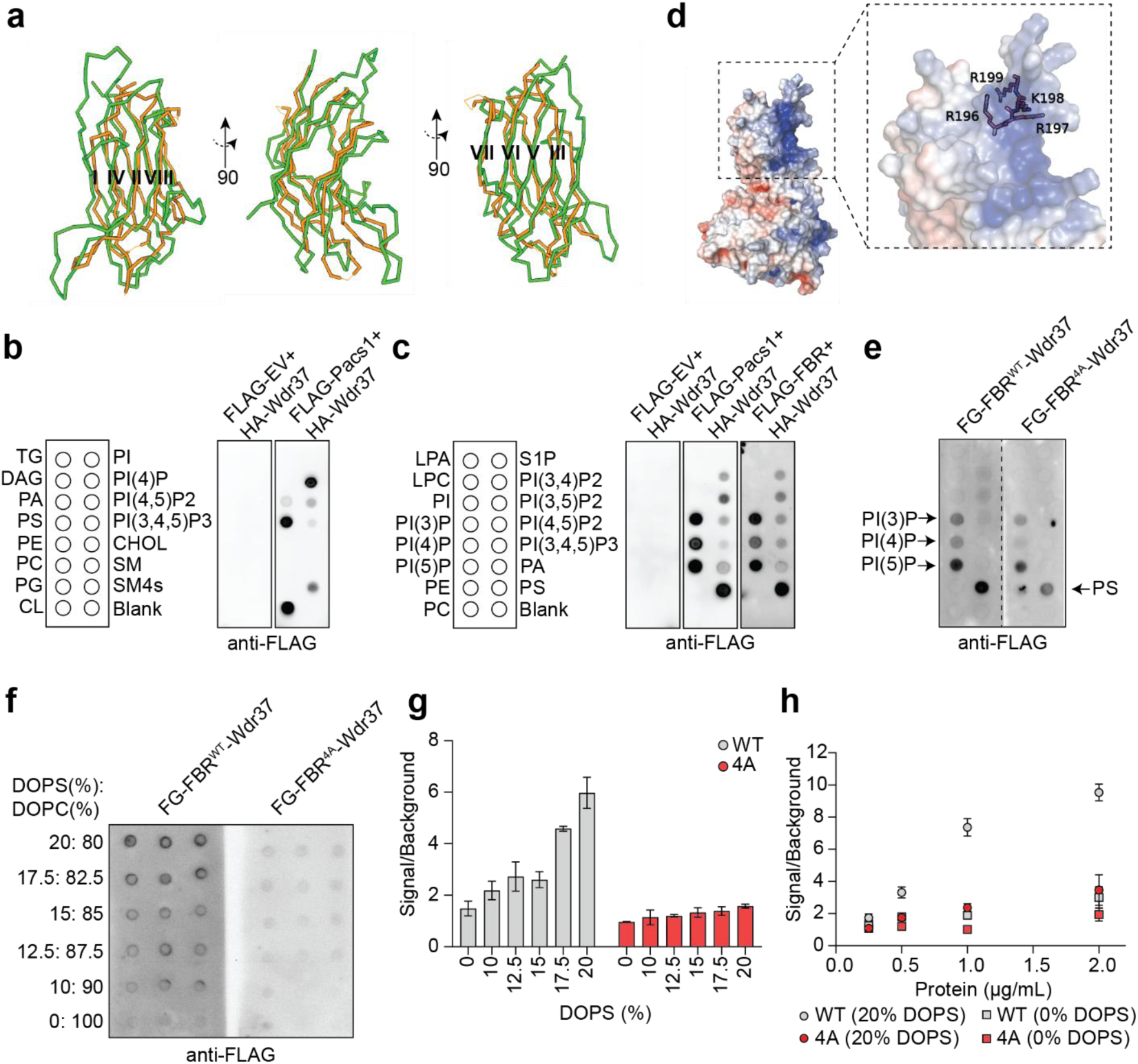
The FBR adopts a lipid-binding fold and binds select acidic phospholipids. **a** Structural overlay of the Pacs1 FBR (green) with the C2 domain of synaptotagmin (orange). **b, c** Protein-lipid overlay assay showing selective binding of Pacs1-Wdr37 complexes to negatively charged membrane lipids. **d** Location of the RRKR motif within the FBR basic cleft. **e** Binding specificity of FG-FBR^WT^ and FG-FBR^4A^ for phosphoinositides (PIPs). **f** Representative dot blot assay to compare binding of FG-FBR^WT^-Wdr37 and FG-FBR^4A^-Wdr37 to liposomes containing indicated ratios of DOPS:DOPC spotted on nitrocellulose membranes. **g** Quantification of liposome dot blots in which 1 µg/ml of FG-FBR^WT^-Wdr37 or FG-FBR^4A^-Wdr37 were incubated with liposome spots containing indicated ratios of DOPS:DOPC. Error bars indicate SD of three technical replicates. Results are representative of two independent experiments. **h** Quantification of dot blots in which indicated concentrations of FBR^WT^-Wdr37 or FBR^4A^-Wdr37 were incubated with membranes spotted with 20% DOPS: 80% DOPC liposomes. Error bars indicate SD of three technical replicates. Results are representative of two independent experiments.

Given the long-standing assumption that Pacs1 functions exclusively through protein-protein interactions, we asked whether the Pacs1-Wdr37 complex might also bind to membrane phospholipids. Using protein-lipid overlay assays, we found that purified Pacs1-Wdr37 complexes bound selectively to negatively charged phospholipids, including phosphatidylserine (PS), cardiolipin, and phosphatidylinositol-4-phosphate (PI4P), while showing no binding to neutral lipids such as phosphatidylcholine (PC) or phosphatidylethanolamine (PE) (Fig. 5b). Further definition of phospholipid specificity showed that the complex also bound to monophosphorylated phosphoinositides PI3P and PI5P but had reduced binding to the poly-phosphorylated PIPs PI(4,5)P₂ and PI(3,4,5)P₃ (Fig. 5c). To test whether this lipid-binding activity was driven by the FBR, we analyzed truncated complexes containing only the FBR and Wdr37. These displayed a similar phospholipid binding profile, indicating that the FBR mediates lipid engagement (Fig. 5C). This binding pattern was similar to that recently reported using purified synaptotagmin C2B domains^19^.

Classical C2 domains bind to phospholipids in a calcium dependent manner^20^. Acidic acid residues within C2 lipid binding loops coordinate calcium ions to increase surface electrostatic charge and promote interaction with negatively charged phospholipids. Notably, the Pacs1 FBR lacks acidic residues in its structurally homologous regions. Instead, the concave face of the FBR β-sandwich is enriched in basic residues which form an electropositive cleft like lipid-binding surfaces found in calcium-independent C2 domains and plekstrin homology (PH) domains^21–23^ (Fig. 5d).

We hypothesized that this basic cleft mediated binding to negatively charged phospholipids. We sought to destroy this pocket by mutating an RRKR motif that comprises its distal surface to alanines (FBR^4A^) (Fig. 5d). We co-expressed human FLAG-GFP-FBR^WT^ (FG-FBR^WT^) or FG-FBR^4A^ together with full-length, strep-tagged human Wdr37 in Expi293F cells. Transfected cells were lysed in a detergent-containing buffer followed by anti-FLAG co-IP. Interestingly, gel filtration of the FG-FBR^4A^-Wdr37 complex from detergent lysates showed a prominent shift toward complex formation compared to FG-FBR^WT^-Wdr37 where a significant amount of FG-FBR^WT^ eluted in later fractions (Fig. S5a, b). Gel-purified complexes containing either FG-FBR^WT^ or FG-FBR^4A^ bound to negatively charged phospholipids on protein-lipid overlay assays in a similar pattern to complexes isolated directly from crude lysates (Fig. 5e).

We next sought to compare the affinity of these FBR variants for liposomes containing different amounts of PS^24^. FG-FBR^WT^ and FG-FBR^4A^ complexes with Wdr37 were purified and incubated with membranes spotted with liposomes that contained different PS:PC ratios. Complexes containing FG-FBR^WT^ bound most strongly to liposomes with 20% PS under calcium-free conditions. There was a significant drop-off in signal when PS was below 17.5% of total lipid (Fig. 5f, g). FG-FBR^4A^ bound 20% PS liposomes ∼3-fold less than FG-FBR^WT^ and did not show significantly more binding compared to PC only liposomes (Fig. 5g, h).

Together, these findings show that the Pacs1 FBR can bind directly to membrane phospholipids in vitro through a highly conserved basic cleft. These results also suggest that Pacs1 may have previously unrecognized direct membrane interaction capabilities in addition to its reported roles in binding the cytosolic domains of transmembrane cargo.

## DISCUSSION

Our study provides the first detailed structural characterization of Pacs1 in complex with Wdr37. We identify the Pacs1 FBR as the critical domain mediating complex formation and find that the binding interface is conserved in Pacs2 and across species. While Pacs1 has historically been viewed as a trafficking adaptor for phosphorylated cargo proteins, our results show that it is also a critical structural scaffold for Wdr37 by stabilizing the WD40 β-propeller and preventing aggregation. Complex formation is essential for the expression and solubility of both proteins which likely reflects a co-translational assembly mechanism.

While many Pacs1 interactors have been identified by co-IP and GST pulldown assays, the structural basis for these interactions has remained undefined. Our findings suggest that a key challenge in understanding Pacs1 function at the structural level is the intrinsic instability of the protein in the absence of Wdr37. Without Wdr37, Pacs1 is poorly expressed and likely misfolded, complicating interpretation of binding studies performed in isolation. The reciprocal stabilization we observe, together with conserved co-dependence across species, indicates that Pacs1-Wdr37 forms a stable core complex that may be required for subsequent client interactions or proper subcellular targeting. Whether binding of the Pacs1-FBR to client proteins or membrane phospholipids allows Wdr37-independent stabilization is an area of active investigation.

The pathogenic R203W mutation in *PACS1* lies within the FBR but does not participate in the interaction with Wdr37. Because complex formation is preserved, expression of the mutant protein depends on Wdr37. Accordingly, targeting Wdr37 using HaloTag-mediated degradation reduced levels of both wild-type and mutant Pacs1 to similar degrees. These findings suggest that pharmacologic degradation of Wdr37 via PROTACs or molecular glues could reduce the accumulation of mutant Pacs1. Notably, recent work has identified the central pores of WD40 β-propellers as promising drug targets, with small molecules shown to engage and modulate the function of WDR proteins such as EED and WDR5 with nanomolar affinity^25, 26^. Given the conservation of the Wdr37 binding interface between Pacs1 and Pacs2, we hypothesize that Wdr37 may serve as a target for therapeutic intervention in PACS-mediated neurodevelopmental diseases.

Current models suggest that Pacs1 and Pacs2 operate in different protein sorting loops within cells. However, Pacs1 and Pacs2 have also been observed to compensate for each other when one is deleted^3, 4^. Moreover, while germline deletion of either Pacs1 or Pacs2 is viable, combined deletion is embryonically lethal^4^. The mechanism behind this compensation is not well understood. Purification of Pacs1-Wdr37 revealed that the complex exists as a tetramer composed of Pacs1-Wdr37 heterodimers joined together through a Wdr37-Wdr37 N-terminal CC interface. Given that Wdr37 interacting residues within the Pacs1 FBR are entirely conserved in Pacs2, it is possible that different combinations of Pacs1 and Pacs2 can bind Wdr37-Wdr37. This scenario would suggest that Pacs1 and Pacs2 are not necessarily separated into different subcellular compartments and that their shared ability to bind Wdr37 allows them to compensate for each other.

The Pacs1 FBR adopts an atypical β-sandwich bearing structural homology to lipid-binding C2 domains. Protein-lipid overlay assays and liposome binding assays confirmed that purified Pacs1-Wdr37 complexes can bind to negatively charged phospholipids. Binding favored phosphatidylserine and, to a lesser extent monophosphorylated phosphoinositides, but did not include polyphosphorylated phosphoinositides. The Pacs1-FBR lacks aspartic acid residues typically found in C2 domains that allow calcium-dependent phospholipid binding^20–22^. Instead, the FBR harbors a basic cleft that promotes calcium-independent phospholipid binding. Disruption of this basic cleft reduced binding to phosphatidylserine-containing liposomes. These findings indicate that Pacs1 may directly engage membrane surfaces with high densities of phosphatidylserine or monophosphorylated phosphoinositides.

While it is still unknown how Pacs1’s functions in cells relate to its ability to bind phospholipids, recent studies have found that Pacs1 orthologs in nematodes and fruit flies localize to the plasma membrane and endomembrane compartments in specific cell types^10, 27^. The ability to directly interact with membranes could help localize Pacs1-Wdr37 to compartments where many of its previously reported protein binding partners reside. Further studies are needed to determine cellular contexts that promote Pacs1 phospholipid engagement such as target membrane phospholipid composition, post-translational modification of Pacs1-Wdr37, or the existence of larger protein complexes that modulate Pacs1 membrane association. Notably, the RRKR basic amino acid motif that we mutated to disrupt the phospholipid binding cleft is also a reported casein kinase 2 binding motif^28^. Furthermore, this FBR basic cleft has features of an acidic cluster binding region^29^. Therefore, it will also be important to determine whether Pacs1-Wdr37 binds client proteins and membrane phospholipids simultaneously, or toggles between these interactions in a regulated or competitive manner.

A limitation of this study is that it does not address the structure or function of the Pacs1 MR and CTR. The MR has been proposed to play an autoregulatory role in controlling the binding of Pacs1 and Pacs2 to client proteins, but we were unable to determine the structural basis for this mechanism. This remains an important area for future investigation, particularly given that pathogenic mutations in the Pacs2 MR cause neurodevelopmental disease. Additionally, while our data show that the N-terminal CC domain of Wdr37 is necessary and sufficient for homodimerization, we did not explore the functional consequences of pathogenic mutations located within the linker region between the CC and the β-propeller. Interestingly, several of these mutations substitute polar or charged residues with hydrophobic or cysteine residues^11, 12^. These could conceivably promote aberrant stabilization of the dimer through hydrophobic interactions or disulfide bonds, respectively. Such a mechanism would be consistent with the gain-of-function effects proposed for the pathogenic variants of *PACS1* and *PACS2*. Furthermore, an important area of future investigation will be to determine how pathogenic mutations in Pacs1, Pacs2, and Wdr37 modulate phospholipid and cargo binding properties of Pacs proteins.

In summary, our findings define the architecture and stability determinants of the Pacs1-Wdr37 complex and uncover new structural features that mediate phospholipid binding. The identification of a conserved, obligate Pacs1-Wdr37 interaction module provides a molecular framework for understanding Pacs protein-mediated trafficking and neurodevelopmental diseases that involve *PACS1*, *PACS2*, and *WDR37*.

## ACKNOWLEDGEMENTS

We thank Joe Gleeson, Yishi Jin, and Taruna Reddy for helpful discussions. This work was funded by NIH R01-AI167920 and a grant from the Pacs1 Foundation (to E.N.-G.), a Ben J. Lipps Research Fellowship from the American Society of Nephrology (to L.X.), an Irvington Postdoctoral Fellowship from the Cancer Research Institute (X.P.), and a subaward from NIH U19-AI100627 (to H.W.).

## AUTHOR CONTRIBUTIONS

Conceptualization (ENG, HW, LX); Data Curation (LX, ENG, HW); Formal Analysis (LX, LC, MG, ENG, HW); Funding Acquisition (ENG, HW); Investigation (LX, LC, MG, MT, NP, SC, ENG, X.P., J.J); Methodology (LX, LC, MG, NP, ENG); Project Administration (ENG, HW); Resources (BB, ENG, HW); Supervision (ENG, HW); Validation (LX, LC, MG, ENG, HW); Visualization (LX, LC, MG, ENG, HW); Writing (original draft) (ENG, LX, HW, MG).

## DATA AVAILABILITY

The datasets generated during and/or analyzed during the current study are available from the corresponding authors on reasonable request.

## MATERIALS AND METHODS

### Plasmids

Human Pacs1-FBR with c-terminal GFP and FLAG tags in pcDNA3.1 and full-length human Wdr37 with c-terminal strep-tag II in pcDNA3.1 were used for protein purification for the cryo-EM studies. For domain truncation analyses and mutagenesis studies, mutations were made as indicated to mouse Pacs1 with an N-terminal FLAG tag within the MSCV vector and to mouse Wdr37 with an N-terminal HA tag within the pCMV3 vector.

### Expression and purification of the Pacs1-FBR and Wdr37 complex

Expi293F cells, maintained in 800 ml of Expi293 Expression Media (ThermoFisher), were grown to 2.0 × 10^6^ cells ml^-1^. These were transiently co-transfected with 0.4 mg of PACS1 plasmid and 0.4 mg of WDR37 plasmid using 2.4 mg of polyethylenimine (Polysciences, Inc.). Cells were fed with 10 mM sodium butyrate and 8 ml of 45% D-(+) glucose solution at 12 h after transfection. Cells were harvested another 48 h later by 20 min centrifugation at 2,500 rpm. The collected cells were lysed in lysis buffer (20 mM Tris pH 8.0, 150 mM NaCl and 1 mM tris(2-carboxyethyl)phosphine hydrochloride) by sonication in the presence of a protease inhibitor cocktail (Roche) and centrifuged at 40,000 rpm for 1 h. The supernatant was collected and incubated with anti-FLAG resin for 1 h at 4 °C, with gentle rotation, and the resin was washed with lysis buffer. Bound proteins were eluted using lysis buffer with the addition of 0.1 mg ml^-1^ FLAG peptide, and concentrated. The concentrated sample was applied to a Superose 6 gel filtration column equilibrated with running buffer (same composition as lysis buffer), and the peak for the Pacs1-Wdr37 was collected and concentrated to 0.4 mg ml^-1^ for EM experiments. Wdr37 alone was expressed and purified similarly.

For lipid binding experiments, Pacs1-Wdr37 complexes were purified following the same procedure except the lysis buffer for Expi293F cells was composed of 20 mM Tris pH 8.0, 150 mM NaCl, and 1% NP-40. Co-immunoprecipitated proteins were changed to a detergent free buffer (20 mM Tris, 150 mM NaCl) during washing of the anti-FLAG resin. Samples were eluted, concentrated, and purified by gel filtration as above. Protein concentrations were measured using bicinchoninic acid assay for downstream experiments.

### Cryo-EM grid preparation and data acquisition

A 3.5 µl drop containing the FBR-WDR37 complex was loaded onto a glow-discharged Quantifoil grid (R1.2/1.3 400-mesh gold-supported holey carbon, Electron Microscopy Sciences), blotted for 3-4 s under 100% humidity at 4 °C, and plunged into liquid ethane using a Mark IV Vitrobot (ThermoFisher). Automated data collection was performed at the Harvard Cryo-EM Center, using SerialEM^30^ on a Titan Krios microscope (ThermoFisher) that operates at 300 keV and is equipped with BioQuantum K3 Imaging Filter (Gatan, slit width 20 eV). Movies were recorded with a K3 Summit direct electron detector (Gatan) operating in counting mode at 105,000 magnification (0.83 Å per pixel). All movies were exposed to a total dose of 54.69 e/Å^2^ over 40 frames at a stage tilt of 37° with a defocus range between −1.0 and −2.0 μm.

### Cryo-EM data processing

All data processing software support was from the SBGrid Consortium^31^. Overall, 1,027 movies were corrected by beam-induced motion using the Relion 3.1 implementation of the MotionCor2 algorithm^32, 33^. The contrast transfer function (CTF) and defocus estimation of micrographs were calculated by CTFFIND4^34^. Images were imported into CryoSPARC^35^ for further data processing. 100 images were selected and auto-picked using the Blob Picker function in CryoSPARC, resulting in 205,296 particles. 2D classification was then used to generate templates for the Template Picker function in CryoSPARC, from which a total of 1,832,228 particles were picked. All particles were extracted at a box size of 256 pixels and then subjected to reference-free 2D classification. Bad particles were rejected, and good 2D classes with clear features were selected.

After rounds of 2D classification, 157,603 particles were selected and applied for ab initio reconstruction with K=3, followed by heterogeneous refinement, resulting in 3 data sets corresponding to 70,488 FBR-WDR37 dimer, 44,580 FBR-WDR37 good monomer and 42,535 bad monomer. Homogeneous refinements were conducted for each data set without applying symmetry, which resulted in a 4.8 Å map for the FBR-WDR37 dimer, and 4.7 Å map for the FBR-WDR37 monomer. For the FBR-WDR37 monomer, non-uniform refinement and local refinement were further performed, resulting in a 4.45 Å map. For the FBR-WDR37 dimer, C2 symmetry were applied during non-uniform refinement and local refinement, leading to 4.46 Å map. AlphaFold3 was used to predict the 1:1 FBR-WDR37 complex structure, as well as the CC dimer of Wdr37. While the former was used as initial model for map interpretations, the latter, which was absent in the map, was used to examine the functional role of the CC dimer. Structures were further refined in Phenix^36^. Atomic models and maps were generated in Chimera^37^.

### Co-immunoprecipitation assays

For co-IP experiments, HEK 293T cells in 6 well plates were transfected with the indicated plasmids using PolyJet transfection reagent. 48h post-transfection cells were lysed in buffer containing 1% NP-40 followed by centrifugation at 12,000 rcf for 10 min at 4 degrees C. Clarified lysates were then incubated with either anti-HA or anti-FLAG antibodies conjugated to magnetic beads for 2h at 4 degrees C, followed by four washes with lysis buffer. Anti-FLAG pulldowns were eluted with 0.15 mg/mL 3x FLAG peptide in TBS. Anti-HA pulldowns were eluted with 0.1 M glycine pH 3. Eluates were analyzed by SDS-PAGE and Western blotting according to standard protocols.

### Protein-lipid overlays

Pacs1-Wdr37 complexes were co-immunoprecipitated from transfected HEK 293T cells with anti-FLAG magnetic beads and eluted with 3x FLAG peptide in TBS. Eluted complexes were diluted in 3 mL of 3% BSA in TBST and incubated with membrane lipid strips (Echelon) for 1h at RT followed by three 5-minute washes in TBST. Lipid strips were then incubated with anti-FLAG antibody followed by washing and incubation with HRP-conjugated secondary antibody. Lipid strips were developed using ECL reagents according to standard Western blot protocols.

### Liposome spotting assays

Liposomes were prepared by mixing DOPS and DOPC stocks (Avanti) in chloroform at the indicated ratios in glass tubes. Lipids were dried under argon followed by overnight drying under vacuum. Lipid films were resuspended in 600 µl of aqueous buffer containing 20 mM Tris and 150 mM NaCl such that the final lipid concentration was 1 mM. Lipid films were allowed to hydrate for 1h at RT with vortexing every five minutes. Hydrated lipids were subjected to five free thaw cycles (1 min liquid nitrogen followed by 3 min RT water bath) and water bath sonication for 30 min. Phospholipid vesicles were then passed 11 times through a 0.1 µm polycarbonate filter to generate liposomes. 3.5 µL of liposomes were spotted in triplicate on nitrocellulose membranes and allowed to dry at RT. Membranes were blocked in TBS with 3% BSA for 1h followed by incubation with indicated amounts of protein for 1h. Subsequent washing and probing was performed the same as for lipid strips except detergent free buffer was used in all steps. For comparison of liposome binding of FBR^WT^ and FBR^4A^, membranes incubated with the same protein concentration were imaged simultaneously. To quantify liposome binding, 8 bit TIF images of developed membranes were analyzed in ImageJ. Images were inverted and background was subtracted using the rolling ball method with a diameter of 20 pixels. Signal within regions of interest containing liposome spots on each membrane were analyzed and normalized to signal within an unspotted background area. Results were presented as signal/background.

---

**Figure S1:**
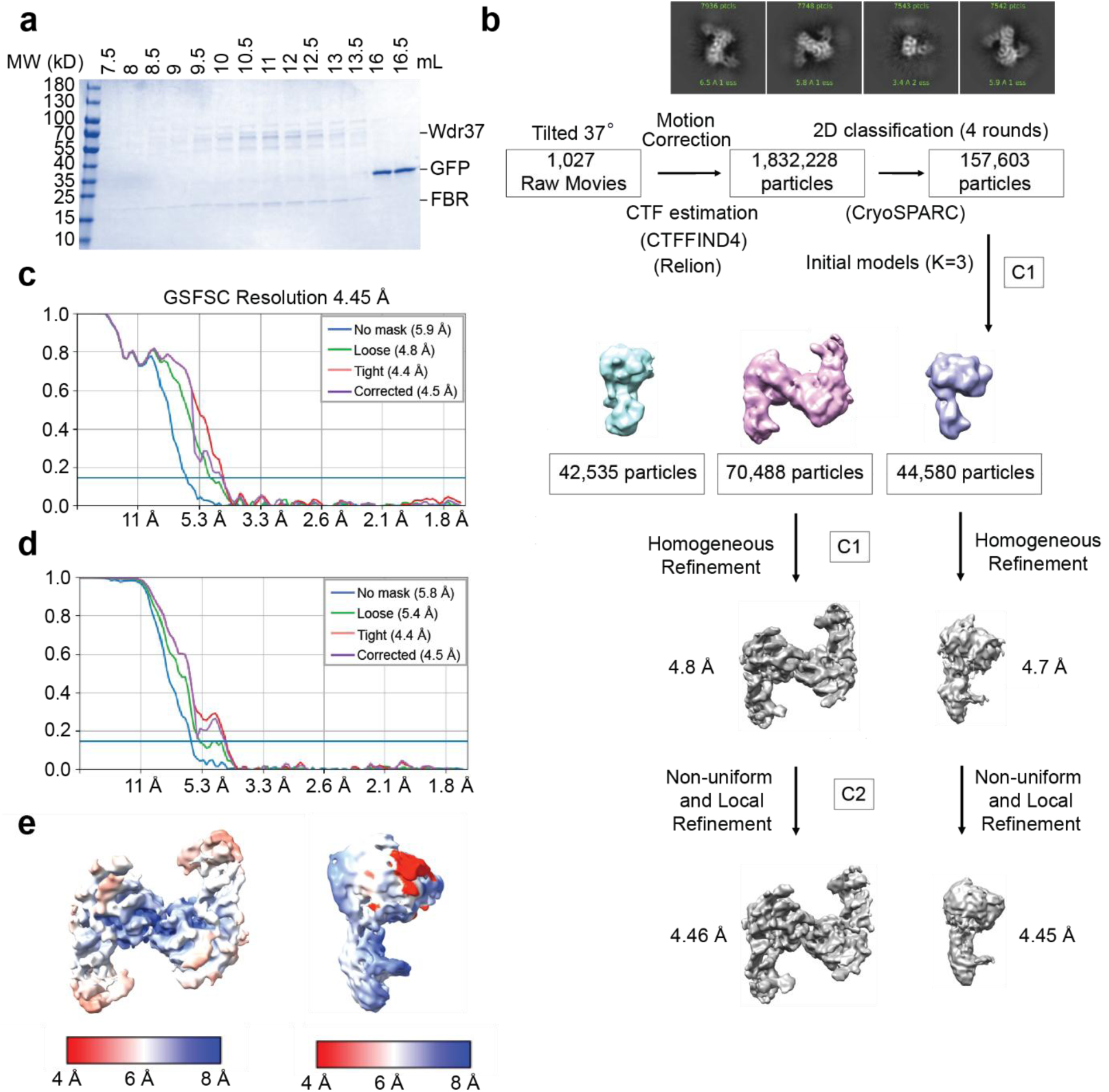
Flow chart of cryo-EM data processing of the FBR-Wdr37 complex. **a** SDS-PAGE of gel filtration fractions from Fig. 1D. **b** Flow chart for cryo-EM processing of the FBR-Wdr37 complex. **c** Fourier shell correlation (FSC) curves for FBR-Wdr37 complex dimer. **d** FSC curves for FBR-Wdr37 complex monomer. **e** Local resolution distribution of FBR-Wdr37 complex dimer. **f** Local resolution distribution of FBR-Wdr37 complex monomer.

**Figure S2:**
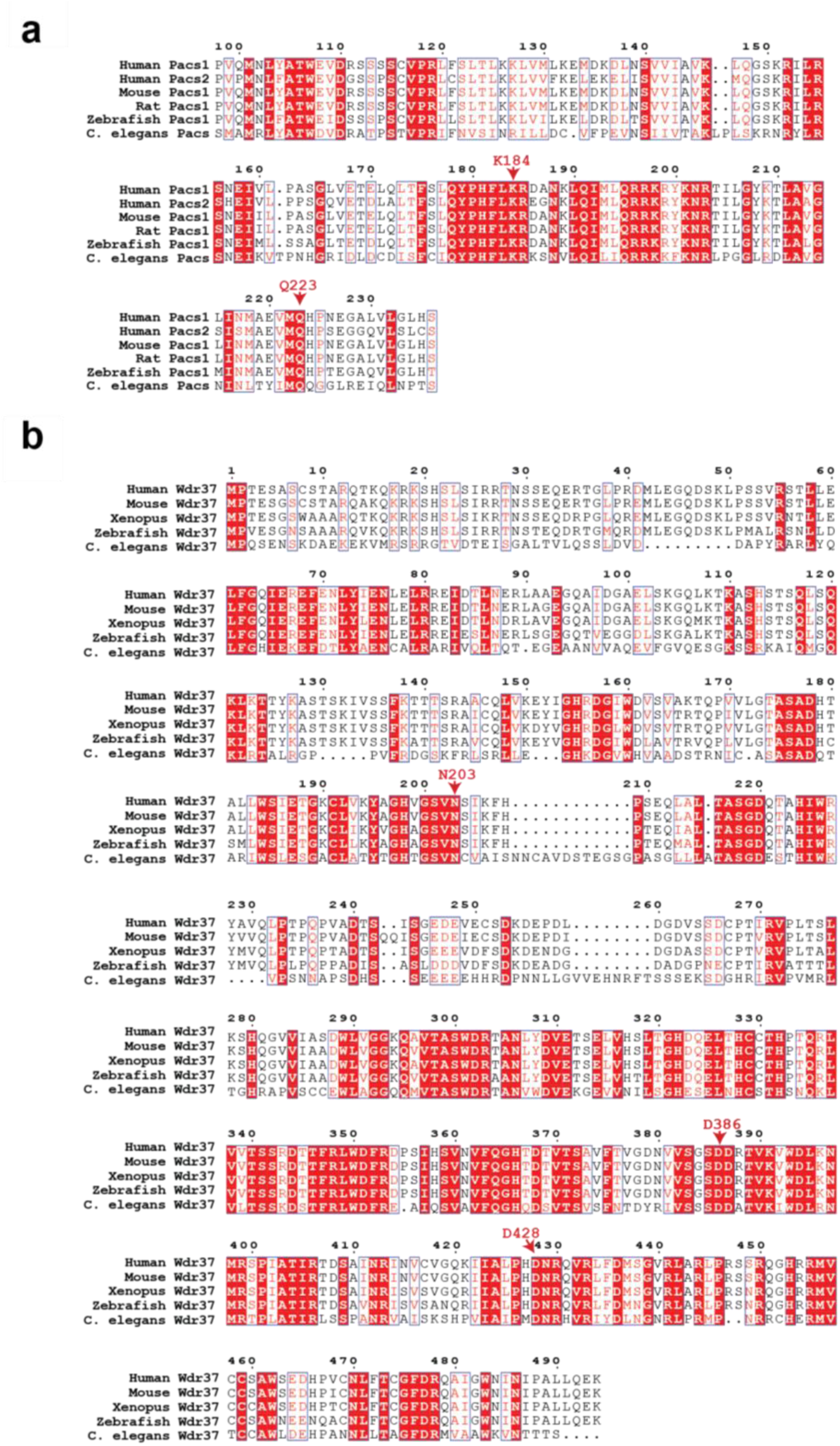
Sequence alignment of the Pacs1-FBR and Wdr37. **a** Multiple sequence alignment of the human Pacs1 FBR with the Pacs2 FBR and orthologs from different species. Wdr37 interacting residues are indicated. **b** Multiple sequence alignment of human Wdr37 with orthologs from different species. Pacs1-interacting residues are indicated.

**Figure S3:**
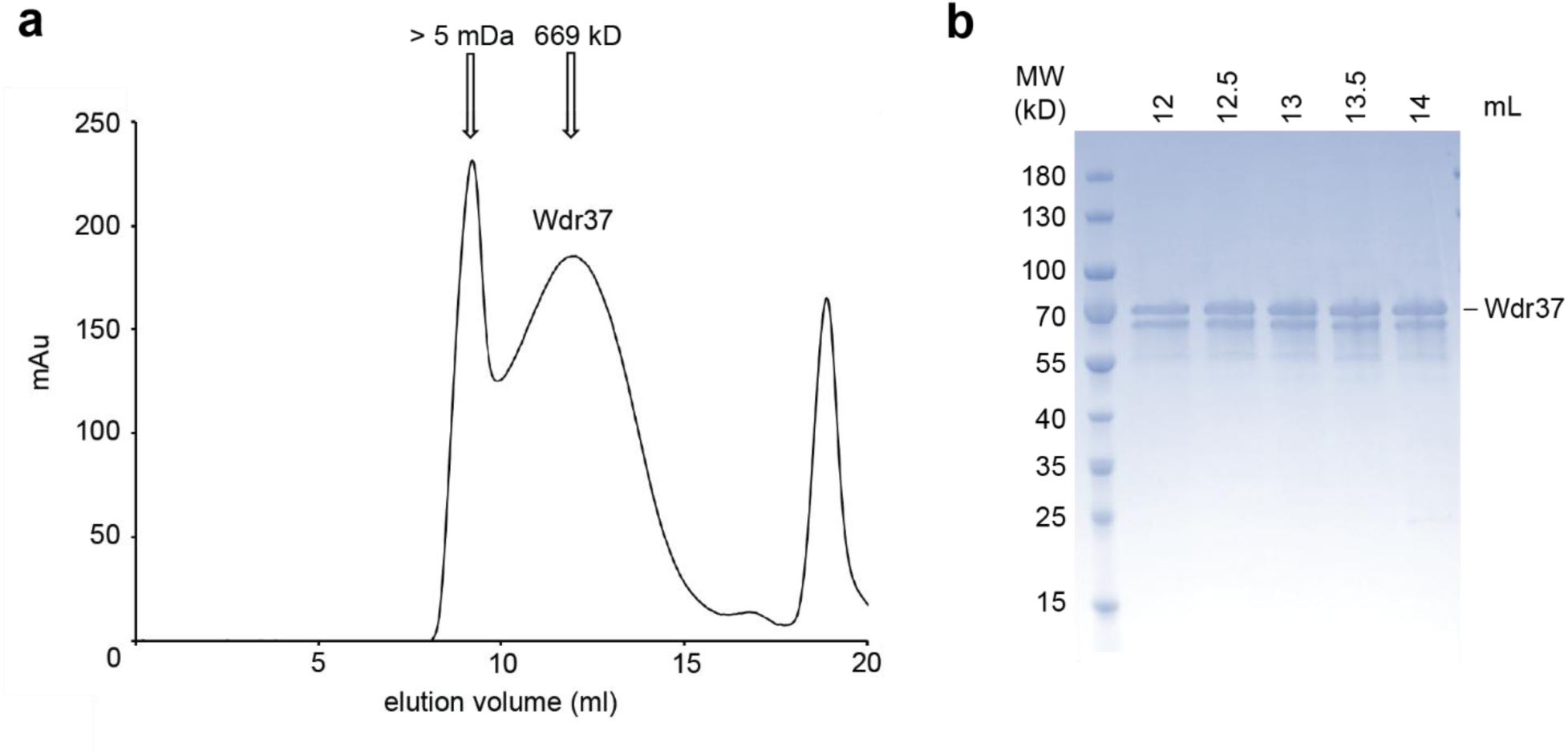
Aggregation of Wdr37 in absence of Pacs1. a, **b** Gel filtration profile and SDS-PAGE analysis showing that expression of Wdr37 without Pacs1 leads to the accumulation of high molecular weight aggregates.

**Figure S4:**
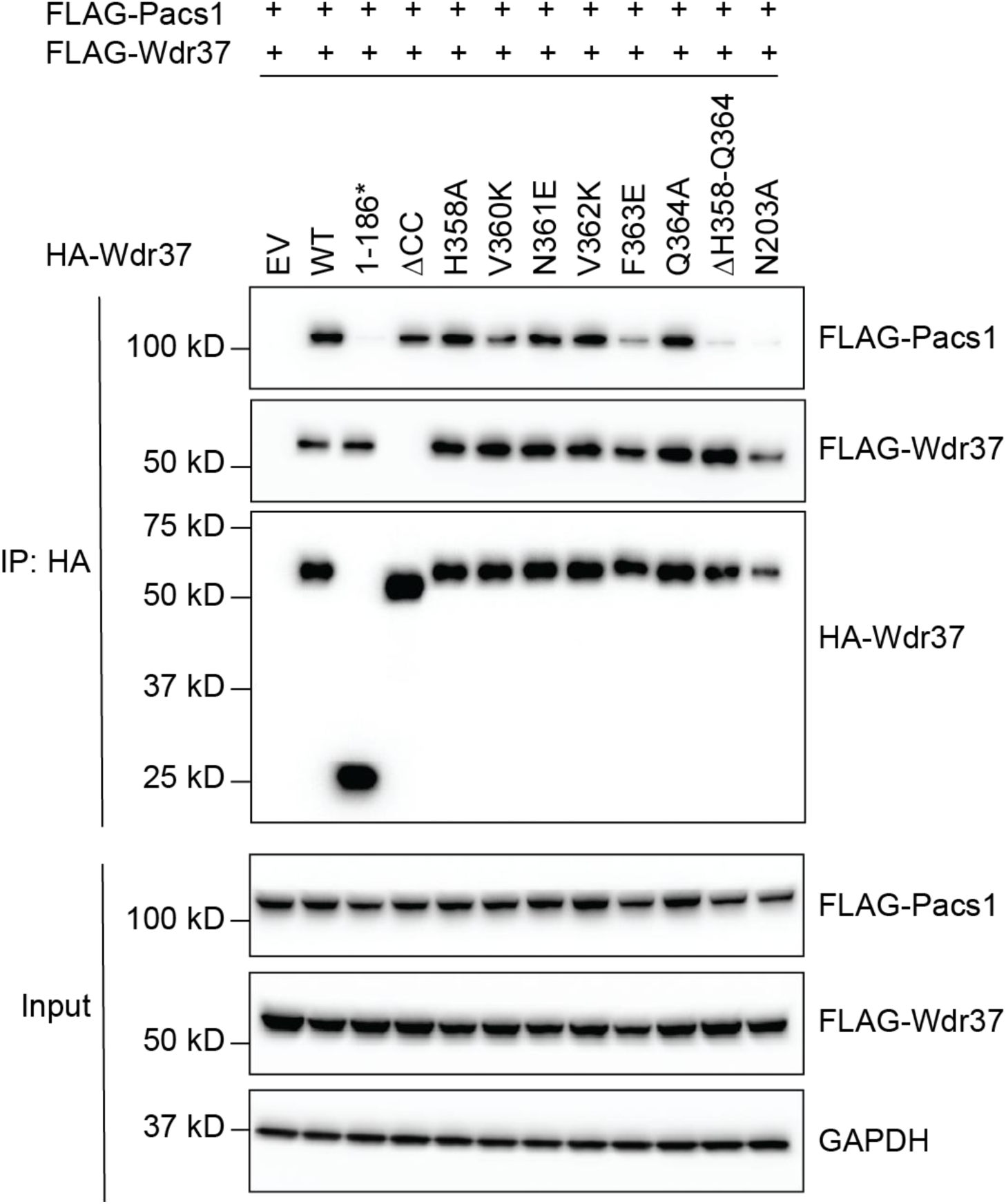
Mutational analysis of the Wdr37 dimer interface within blade 4. Point mutations in HA-Wdr37 were made to disrupt hydrophobic interactions between blades 4 of the Wdr37 homodimer. Co-IP of HA-tagged Wdr37 with WT FLAG-Wdr37 and WT FLAG-Pacs1 was performed using anti-HA beads.

**Figure S5:**
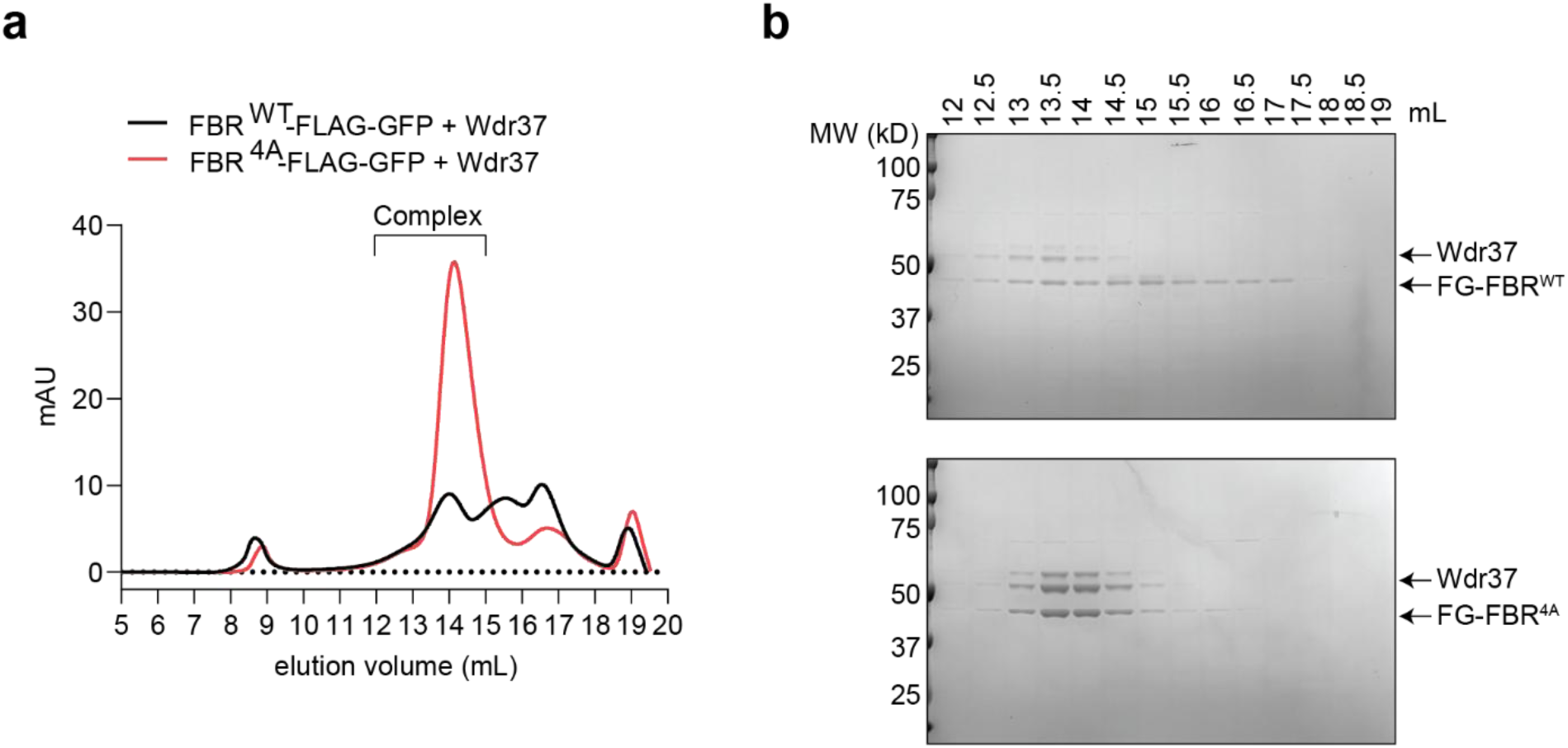
Purification of FG-FBR^WT^-Wdr37 and FG-FBR^4A^-Wdr37 complexes from detergent lysates. **a** Gel filtration profiles of FG-FBR^WT^-Wdr37 and FG-FBR^4A^-Wdr37 complexes. **b** SDS-PAGE of fractions collected during the gel filtration of FBR WT and 4A.

**Table S1.** Cryo-EM data collection, refinement, and validation statistics.

|  | FBR-Wdr37 (dimer) | FBR-Wdr37 (monomer) |
| --- | --- | --- |
| **Data collection and processing** |  |  |
| Nominal Magnification | 105,000 | 105,000 |
| Voltage (keV) | 300 | 300 |
| Electron exposure (e<sup>-</sup>/Å<sup>2</sup>) | 54.69 | 54.69 |
| Defocus range (μm) | -1.0 – -2.0 | -1.0 – -2.0 |
| Pixel size (Å) | 0.825 | 0.825 |
| Symmetry imposed | C2 | C1 |
| Initial particle images (no.) | 1,832,228 | 1,832,228 |
| Final particle images (no.) | 70,488 | 44,580 |
| Map resolution (Å) | 4.46 | 4.45 |
| FSC threshold | 0.143 | 0.143 |
| Map resolution range (Å) | 4.0 – 6.0 | 4.0 – 6.0 |
| **Refinement** |  |  |
| Initial model used | AlphaFold | AlphaFold |
| Model resolution (Å) | 4.4 | 4.4 |
| FSC threshold | 0.143 | 0.143 |
| Map sharpening *B* factor (Å<sup>2</sup>) | -287.8 | -328.8 |
| Model composition |  |  |
| Non-hydrogen atoms | 7,293 | 3,721 |
| Protein residues | 933 | 476 |
| Ligands | - | - |
| ***B*-factors(Å<sup>2</sup>)** |  |  |
| Protein | 163.79 | 84.79 |
| Ligand | - | - |
| **R.m.s. deviations** |  |  |
| Bond lengths (Å) | 0.005 | 0.002 |
| Bond angles (°) | 0.832 | 0.649 |
| Validation |  |  |
| MolProbity score | 2.24 | 1.97 |
| Clashscore | 22.63 | 16.84 |
| Poor rotamers (%) | 0 | 0 |
| **Ramachandran Plot** |  |  |
| Favored (%) | 94.25 | 96.38 |
| Allowed (%) | 5.54 | 3.62 |
| Disallowed (%) | 0.22 | 0 |

